# FGF12A Regulates Nav1.5 via CaM-regulated and CaM-independent Mechanisms

**DOI:** 10.1101/2025.01.02.631125

**Authors:** Lucy Woodbury, Paweorn Angsutararux, Martina Marras, Emily Wagner, Carlota Abella, Anna Li, Jonathan R. Silva

**Affiliations:** Department of Biomedical Engineering, McKelvey School of Engineering, Washington University in St. Louis, St. Louis, MO, USA; Department of Pharmacology, School of Medicine, UC Davis, Davis, CA, USA; Baylor College of Medicine, Houston, TX, USA

## Abstract

Opening of the cardiac voltage-gated Na+ channel (Nav1.5) is responsible for robust depolarization of the cardiac action potential, while inactivation, which rapidly follows, allows for repolarization. Regulation of both the voltage- and time-dependent kinetics of Nav1.5 inactivation can alter the ability of the heart to initiate and sustain a re-entrant arrhythmia. The C-terminal domain (CTD) of Nav1.5 has been shown to modulate fast inactivation of the channel, and multiple auxiliary proteins bind to the CTD, including calmodulin (CaM) and intracellular fibroblast growth factor 12A (FGF12A). Recently, a non-canonical CaM-binding site was also discovered on the N-terminal of A-splice variants of iFGFs. We performed cut-open Vaseline gap (COVG) voltage-clamp to test whether FGF12A with and without CaM regulates Nav1.5 gating. In WT Nav1.5 channels, FGF12A with and without CaM present had a minimal effect on the voltage dependence of both activation and inactivation. Conversely, when CaM is absent on the Nav1.5 CTD (IQ/AA), a dramatic shift in steady-state inactivation (SSI) occurred, regardless of whether CaM was present on FGF12A. These two distinct mechanisms are operative in Nav1.5 LQT3 mutations where FGF12A requires CaM to shift in the voltage-dependence of inactivation, but not to inhibit the persistent late current. We conclude that there are two distinct mechanisms by which FGF12A modulates the Nav1.5 channel: CaM-regulated alteration of the voltage dependence of inactivation and CaM-independent inhibition of persistent late current.

## Introduction

Voltage-gated sodium (Nav) channels are responsible for the initiation of the action potential in excitable cells including neurons and myocytes. The Nav1.5 sodium channel, encoded by the *SCN5A* gene, is the most prevalent Nav channel in cardiomyocytes and is responsible for the rapid post-stimulus depolarization of the myocyte (Catterall et al., 2005). Nav1.5 contains four homologous repeats (I-IV) each comprising six transmembrane helices (S1-S6) with the first four defining the voltage-sensing domain (VSD). The S4 helix carries multiple positive charges, which translocate in response to depolarization of the cell membrane to actuate channel gating (Catterall, 2023; Catterall et al., 2005; D et al., 2020). Nav1.5 gating has been described in terms of closed (deactivated), open (activated), and inactivated states (Ahern et al., 2016; Mangold et al., 2017) where channels transition from closed to open in response to depolarizing changes in membrane potential and then to inactivated as an elevated membrane potential is maintained. From the closed state, strong depolarization of the plasma membrane will activate the first three VSDs (I-III). Notably, VSD-IV is not required for channel-pore opening, however it is required for inactivation (Capes et al., 2013; Chanda and Bezanilla, 2002). The activation of the VSD-IV allows a motif in the intracellular III-IV linker, isoleucine-phenylalanine-methionine (IFM), to bind to a hydrophobic pocket that is proximal to the intracellular side of the pore (D et al., 2020; Jiang et al., 2021). The IFM motif then allosterically alters the conformation of the S5-S6 helices to close the pore and produce a fast inactivation of the channel, allowing the cardiac action potential to eventually repolarize, returning the membrane to negative potentials (Li et al., 2021). The C-terminal domain (CTD) of Nav1.5 has also been implicated in modulating activation and inactivation kinetics (Deschênes et al., 2001). Recent structures of Nav1.5 channel in its open state have shown the channel CTD interacting with the III-IV linker, modulating fast inactivation (Biswas et al., 2024; Shen et al., 2017; Wang et al., 2012).

Numerous variants within *SCN5A* are known to predispose patients to cardiac arrhythmia. This pathology can arise when variants alter the peak and late components of the Na^+^ current (I_Na_) causing either a gain or loss of function. One of these, Long QT syndrome (LQTS), arises from the prolongation of ventricular repolarization that manifests as a prolonged ventricular myocyte action potential and corrected Q-T interval of an ECG. LQTS can be caused by gain-of-function mutations that typically affect the late or persistent Na^+^ current (I_Na-L_). Normally, I_Na,L_ is relatively small compared to the peak I_Na_ (0.1 – 1%). Variants that significantly increase I_Na,L_ bring excess positive charges into the myocyte that prolong the action potential, leading to proarrhythmic triggered activity that includes early and delayed afterdepolarizations (Han et al., 2018). Loss-of-function mutations within SCN5A are usually associated with Brudaga syndrome which has over 300 associated variants (Mizusawa and Wilde, 2012), which can cause slowed conduction and dispersion of repolarization.

Multiple subunits interact with Nav1.5 to regulate its function and pharmacology (Abriel, 2010). The β1 (*SCN1B*) subunit alters the voltage dependence of DIII and DIV VSD activation, Nav1.5 gating kinetics, and anti-arrhythmic drug affinity (Angsutararux et al., 2021; Martinez-Moreno et al., 2020; Moreno et al., 2019; Zhu et al., 2021, 2017). Calmodulin (CaM) alters trafficking and modulates the voltage sensing kinetics of multiple channels including KCNQ1, Nav1.5, and Cav1-2 (Ben-Johny and Yue, 2014; Crotti et al., 2013; Herzog et al., 2003; Houdusse et al., 1996; Kang et al., 2023; Yu et al., 2003). CaM itself binds to the IQ (Isoleucine-Glutamine) motif located at Nav 1908-09 on the Nav1.5 CTD (Kim et al., 2004; Tan et al., 2002). The CaM protein itself is a ubiquitous Ca^2+^ sensor that is made up of four E-F hands arranged in a dumbbell-like manner. CaM will change its conformation or binding orientation based on the addition of Ca^2+^ on its C and N-lobes. Typically, CaM must be saturated with Ca^2+^ at all four E-F hands to bind to various proteins including KCNQ1 and FGF12A (Mahling et al., 2021; Shamgar et al., 2006). However, apo-CaM without Ca^2+^ can interact with the Nav1.5 CTD (Ben-Johny et al., 2016). The loss of CaM binding is associated with LQTS, Brugada Syndrome, and atrial fibrillation (Kapplinger et al., 2010; Wilde and Amin, 2018). CaM binding to Nav1.5 is seen as protective against increased persistent late current and allows for increased peak channel opening (Kang et al., 2021; Yan et al., 2017).

Intracellular fibroblast growth factors (iFGFs), also known as fibroblast homologous factors (FHFs), have distinct isoforms with varying exon splicing (e.g., FGF12A and FGF12B). These iFGFs do not have the N-terminal sequence that is required for extracellular secretion and therefore do not bind to FGF receptors (Pablo and Pitt, 2016). However, specific iFGFs do modulate voltage-gated ion channels including sodium and calcium channels (Pablo and Pitt, 2016). iFGFs interact with the Nav1.5 channel through the iFGF β-trefoil at the EF-hand of the Nav1.5 (Goetz et al., 2009; Goldfarb, 2024). The interaction between FGF12 and Nav1.5 is known to induce a depolarizing shift in the steady-state inactivation (SSI) of the channel, leading to a reduction in the number of channels inactivated at a resting membrane potential (Angsutararux et al., 2023; Dover et al., 2010; Goldfarb, 2024). FGF12A specifically has also been shown to reduce pathogenic persistent late sodium current (I_Na,L_) carried by Nav1.5 channels (Chakouri et al., 2022). The Nav1.5 variant of H1849R blocks binding of iFGFs to the Nav1.5 channel which resulted in a gain-of-function increase of I_Na_ by altering steady-state inactivation and ultimately resulting in arrythmia (Musa et al., 2015). FGF12B and FGF13VY have also been shown to alter the decay phase of peak I_Na_ after fitting the curves to the sum of two exponentials which are on the time scales of 1ms (T_1_) and 10ms (T_10_) time components of inactivation (Angsutararux et al., 2023).

All A-splice variants of iFGFs have recently been shown to have a non-canonical CaM binding site on the N-terminal portion of the protein (Mahling et al., 2021). The CaM binding region on FGF12A allows for the possibility of multiple calmodulins binding to the Nav1.5 CTD. The relationship between CaM and FGF12A independently on the CTD has been analyzed (Gade et al., 2019), however, the specific interactions of the CaM located on the FGF12A N-terminus with Nav1.5 gating kinetics has not been defined. We have designed experiments to test whether CaM presence on FGF12A is required for the myriad changes in Nav1.5 kinetics that have been previously reported, including shifts in activation, inactivation and reduction of I_Na,L_ (Chakouri et al., 2022).

## Materials and Methods

### *Xenopus Laevis* oocyte collection

*Xenopus laevis* females with length > 9cm were purchased from Xenopus One Corp. Oocytes were collected from anesthetized frogs in accordance of previously reported protocols(Varga et al., 2015). All animal handling protocols were approved by Washington University’s Institutional Animal Care and Use Committee (IACUC) and were developed in compliance with the National institutes of Health (NIH) Guild for the Care and Use of Laboratory Animals.

### Molecular biology

The molecular biology protocols used were adapted from Angsutararux et al. 2023. Plasmids encoding human *SCN5A* and *Fgf12A* were produced on a pMAX and pBSTA backbone respectively. All Nav1.5 point mutations (**Table 1**) and FGF12A del CaM (**Figure 1**) were generated using High-Fidelity PCR via the Herculase II Fusion DNA Polymerase (Agilent). The scrambled FGF12A (FGF12A Scram CaM) sequence was created with GenScript (**Figure 1**). The plasmid was constructed and packaged by VectorBuilder with the vector ID of VB231023-1405afd. cDNA plasmids were amplified from glycerol E. coli stocks with NucleoSpin Plasmid DNA purification (Macherey-Nagel). The cDNA plasmids were linearized with the NucleoSpin Gel and PCR Clean-up kit (Macherey-Nagel). cRNA was then synthesized with the mMessage mMACHINE T7 Transcription kit (Life Technologies) and reconstituted in nuclease-free H_2_O at a concentration of approximately 1 μg/μL.

**Table 1.**
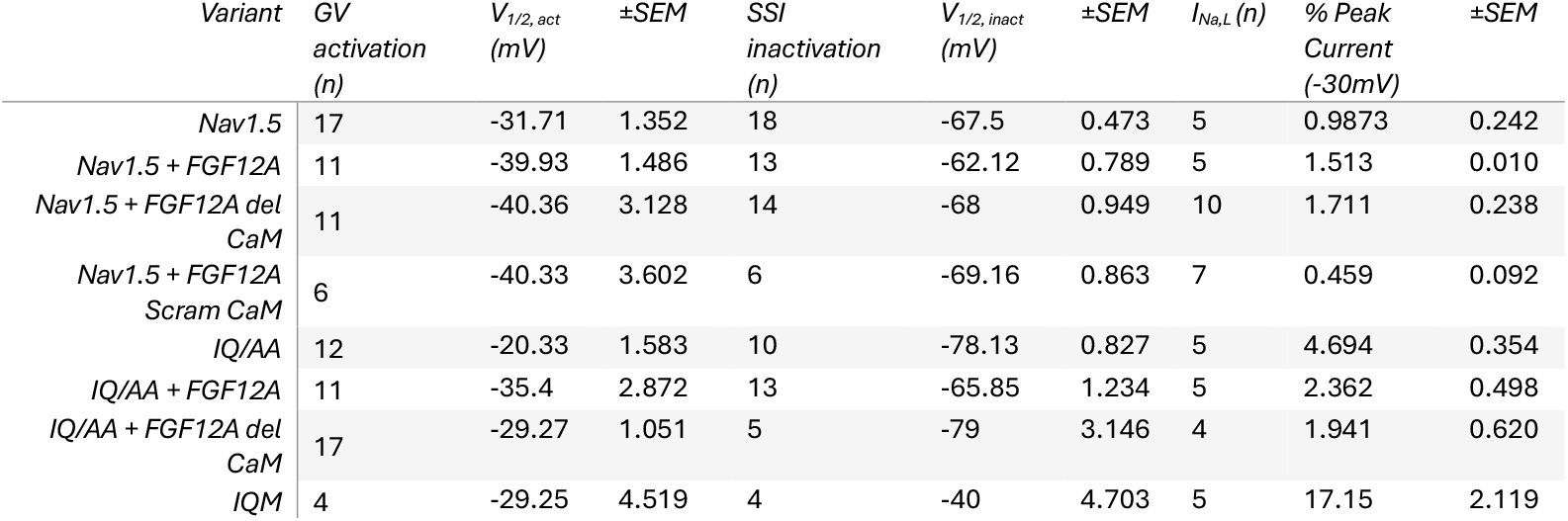
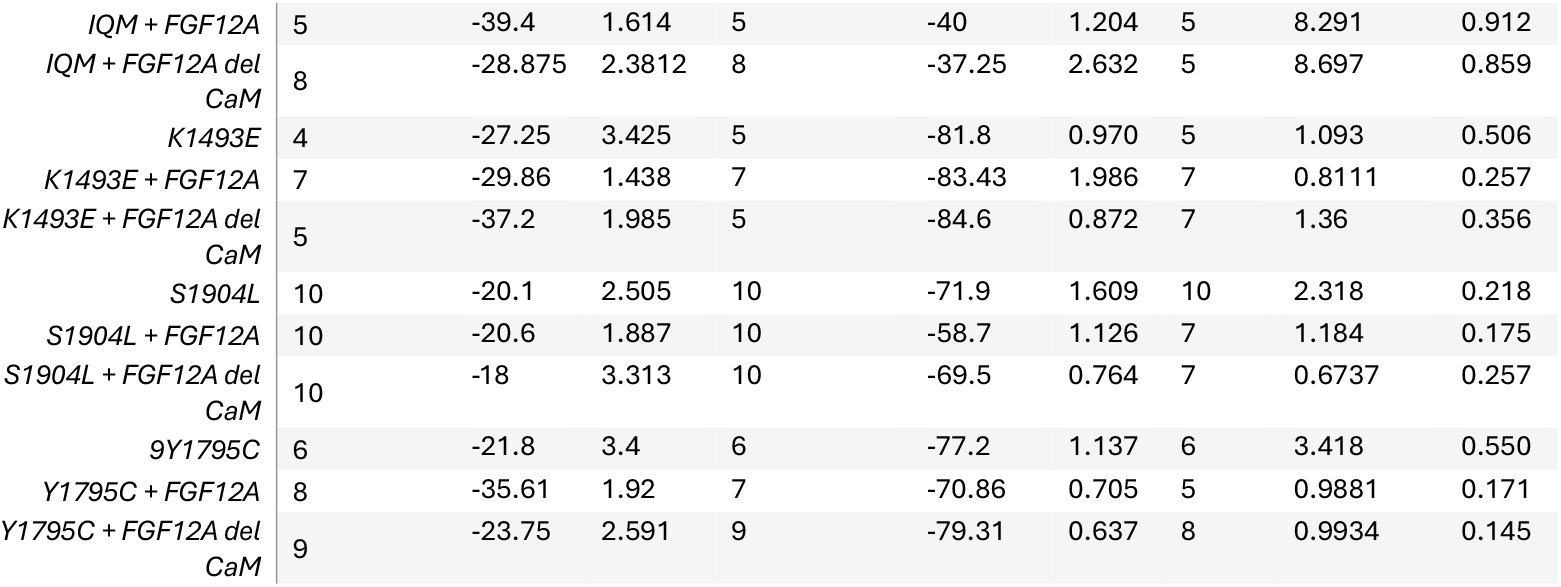

**Figure 1:**
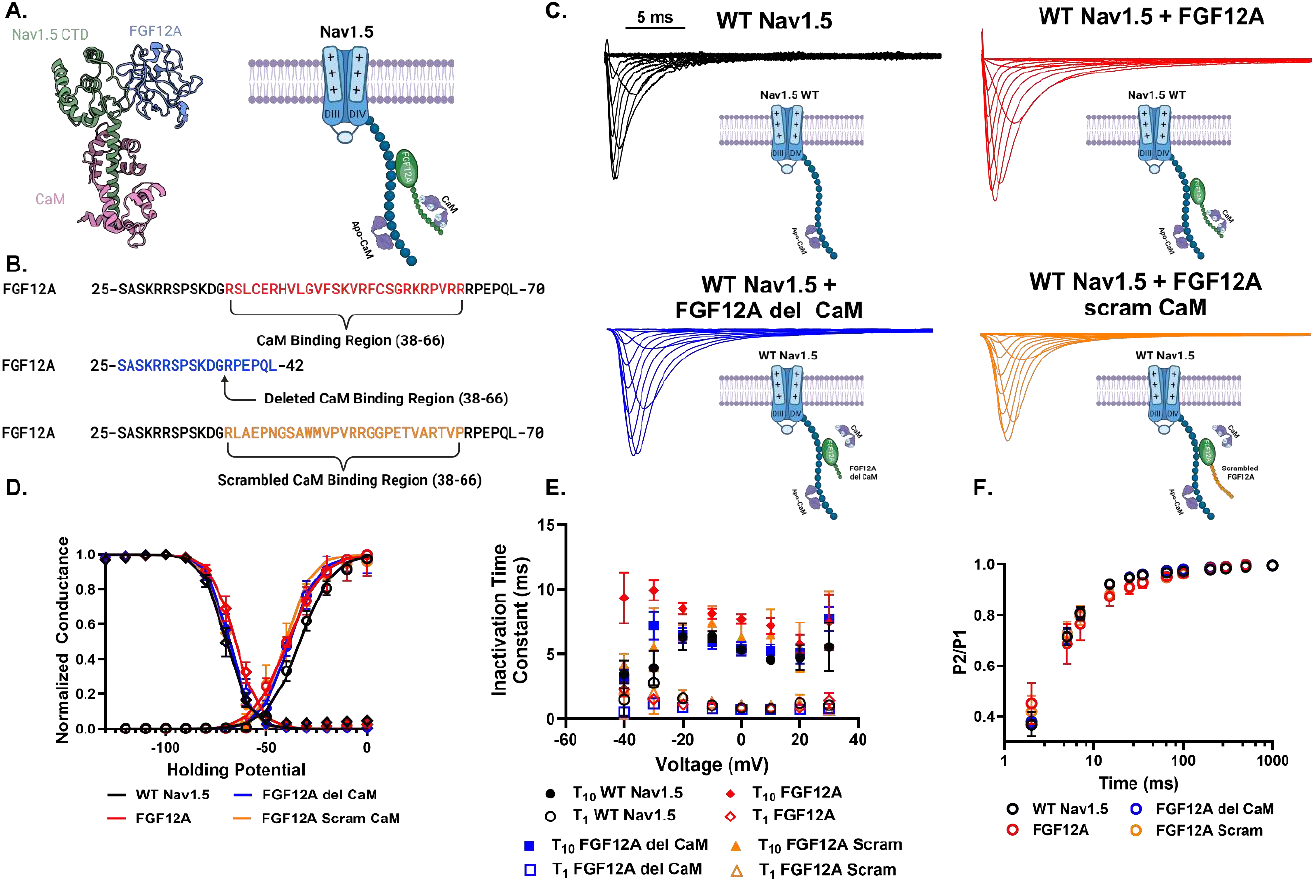
CaM binds to FGF12A non-canonically to confer shifts in inactvation kinetics. **A)** Schematic representation of the Nav1.5 C-terminus domain (CTD) with auxiliary proteins calmodulin (CaM) and intracellular fibroblast growth factor 12A (iFGF12A), not drawn to scale. **B)** Sequence of CaM binding region describe in Mahling et al. 2021. Red denotes WT FGF12A, Blue: deleted CaM binding region (FGF12A del CaM), Orange: scrambled CaM binding region (FGF12A Scram CaM). **C)** Representative traces with corresponding auxiliary subunit schematics. **D)** Mean normalized ±SEM conductance-voltage (GV) and steady-state inactivation (SSI) curves for WT Nav1.5 and the addition of: FGF12A, FGF12A del CaM, and FGF12A Scram CaM. Curves are fitted single Boltzmann functions. The absence of CaM on FGF12A shows a slight shift in the inactivation of the Nav1.5 (**Table 1**). **E)** Mean ±SEM T1 and T_10_ inactivation time constants of I_Na_ decay fitted to a double exponential described in Methods. WT FGF12A showed a marked increase in the slow time component at multiple holding potentials. **F)** Mean ±SEM recovery from inactivation from a protocol described in Methods.

### Cut-open oocyte voltage-clamp

Collected *Xenopus laevis* were digested with 24mg collagenase and 3mg trypsin inhibitor in 35 mL OR2 buffer (in mM, 82.5 NaCl, 2.5 KCl, 1 MgCl_2_, 5 HEPES, pH 7.6) for 20 minutes. Oocytes were selected and kept at 5°C overnight in ND93 solution (in mM, 93 NaCl, 5 KCl, 1.8 CaCl_2_, 1 MgCl_2_, 5 HEPES, 2.5 Na pyruvate, and 1% penicillin/streptomycin, pH 7.4). Oocytes were injected with 50ng of cRNA with a ratio of 2:1:1 (Nav1.5, β_1_, FGF12A) for co-expression or 3:1 (Nav1.5:β_1_) and incubated for 24 hours at 18°C prior to recordings. The auxiliary subunit of β_1_ (*SCN1B*) was present in each group for consistency and increased channel expression (Martinez-Moreno et al., 2020).

A custom cut-open Vaseline gap (COVG) rig and protocol described previously was used for oocyte voltage-clamp recordings (Angsutararux et al., 2023; Rudokas et al., 2014; Varga et al., 2015). Oocytes were kept at 18 C with an HCC-100A temperature controller (Dagan Corporation) throughout recordings. A CA-1B cut-open amplifier (Dagan Corporation) with corresponding electrodes and bath-guard connected with a digitizer (1440 Digidata; Molecular Devices) to the computer software Clampex (Molecular Devices). The oocytes were recorded in an external solution (in mM, 25 N-methyl-D-glucamine (NMDG), 90 2-(N-Morpholino)ethanesulfonic acid (MES) sodium salt (Na-MES), 20 HEPES, 2 Ca-MES_2_, pH 7.4) and an internal solution (in mM, 105 NMDG, 10 Na-MES, 20 HEPES, 2 EGTA, pH 7.4).

For steady-state activation, oocytes were clamped from -120 to 60mV for 100ms with a holding potential of -120mV. The peak currents were measured, normalized, and divided by the driving force to determine the conductance (G) at each voltage step. The G-V curves were fitted with the Boltzmann Equation (Eq. 1):

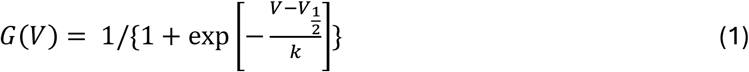

with V_1/2_ as the voltage at half-activation and k is the slope. To measure the inactivation time components, a two-exponential equation (Eq. 2) was fitted to the decay phase of activation at each voltage step:

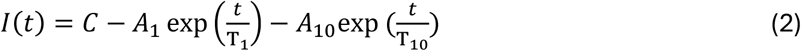

where A_1_ and A_10_ are the amplitudes of inactivaiton and T_1_ and T_10_ are the time components on the time scale of 1ms or 10ms respectively.

Steady-state inactivation involved pre-pulsing the cells from -150 to 20mV for 200ms and then stepping to - 20mV to determine the channel availability. Recovery from inactivation was a two-pulse protocol in which the first pulse stepped up from the holding potential of -120mV to -20mV for 20ms. The second pulse was applied after a varying amount of time (2-1000ms). The ratio between the first peak (P1) and the second (P2) was used to determine channel recovery.

For intermediate inactivation prolonged protocols were used during voltage-clamp experiments(Veldkamp et al., 2000). Intermediate inactivation followed a similar protocol to steady-state inactivation except with a 1000ms depolarizing pulse before measuring channel availability a -20mV. Intermediate recovery was similar to the recovery from inactivation protocol except the first pulse had a 1000ms duration. Intermediate onset included a two-pulse protocol where the first pulse was a step to -20mV for a duration varying from 2-1000ms. A second pulse was applied after 20ms to determine the time-course of intermediate inactivation (Wang et al., 2000).

### Electrophysiological data analyses

Electrophysiological data and statistical analysis were conducted with Clampfit (Molecular Devices), Prism 10 (Graph Pad), and JupyterLab. The statistical significance between experimental groups was determined by a Welch ANOVA unpaired t-test or Tukey’s multiple comparisons test.

### Supplemental material

**SI Figure 1** includes the conductance-voltage relationship curve (GV) and steady-state inactivation curves (SSI) for the Nav1.5 variants of Y1795C, S1904L, K1493E, and IQM compared to WT Nav1.5 and co-transfected with either WT FGF12A or FGF12A del CaM. **SI Figure 2** shows the recovery from inactivation curves of the Y1795C, S1904L, and K1493E Nav1.5 variants with either WT FGF12A or FGF12A del CaM. **SI Figure 3** are the inactivation time constants fitted from the decay phase of the activation pulse protocol for the aforementioned Nav1.5 variants with either WT FGF12A or FGF12A del CaM. **SI Table 1** includes all of the fitted inactivation time constants at - 30mv for all constructs included in the project. SI Figure 4 shows the representative traces of Y1795C, S1904L, and K1493E co-transfected either with WT FGF12A or FGF12A del CaM.

## Results

### A-splice variants of iFGFs have a non-canonical CaM binding region

We tested whether the presence of CaM on FGF12A affects FGF12A regulation of Nav1.5. To prevent CaM interaction with FGF12A, we considered two options: deleting the entire CaM binding region (*residues 38-66*) on FGF12A (FGF12A del CaM) or scrambling the CaM binding region section of FGF12A (FGF12A scram CaM) (**Figure 1B**). We conducted cut-open voltage clamp recordings with Nav1.5 expressed alone or with: FGF12A (*red*), FGF12A del CaM (*blue*), and FGF12A scram CaM (*orange*) (**Figure 1C**). Co-expression of WT FGF12A resulted in a small depolarizing shift in the SSI (+5.385mV ±1.546, p = 0.0462) and a hyperpolarizing shift in GV curve (−8.226mV ± 2.065, p=0.0005) (**Figure 1D and Table 1**), consistent with previous observations (Dover et al., 2010). Inactivation was well-fit with two exponentials with time constants on the order of 10ms and 2ms. The 10ms inactivation time constant (T_10_) was significantly increased with WT FGF12A co-expression(+2.98ms ±0.667, p = 0.029, **Figure 1E**). Conversely, with the CaM binding region deleted from FGF12A, no shifts in T_10_ (−0.729ms ±0.462, p=0.1585) (**SI Table 1**) or SSI V_1/2_ (−0.5mV ±0.992, p=0.6179) (**Table 1**) were observed.

We continued by comparing FGF12A scram CaM with FGF12A del CaM, as the deleted section of the FGF12A protein was substantial. No significant changes in FGF12A scram CaM’s effects compared to the FGF12A del CaM protein was detected (**Figure 1C-F and Table 1**). Therefore, we concluded that the depolarizing shift in SSI voltage-dependence and inactivation time constants were due to the lack of CaM binding itself and not changes in overall protein structure of FGF12A that were caused by the deletion of the CaM binding region. Subsequent studies used FGF12A del CaM as the sole source of CaM deletion from FGF12A.

### CaM’s absence from the Nav1.5 CTD allows for an increased effect due to FGF12A

Results from **Figure 1** show that FGF12A can only shift WT Nav1.5 inactivation when CaM is bound to the FGF12A N-terminus. Therefore, we considered whether the removal of CaM from the Nav1.5 CTD would amplify the FGF12A effect, given that the only CaM present in the Nav1.5 complex would be associated with WT FGF12A. We proceeded to test this hypothesis by using the IQ(1908-09)/AA variant that inhibits CaM binding to Nav1.5 (Abrams et al., 2020) (**Figure 2A**). The IQ/AA variant of Nav1.5 independently has large hyperpolarizing and depolarizing shifts in its SSI (−10.63mV ±1.676, p<0.0001) and GV V_1/2_ (+11.37mV ±2.65, p=0.003) respectively and significantly increases the persistent late current (I_Na,L_) (+0.037% ±0.006, p<0.0001) compared to WT Nav1.5 (**Table 1**).

**Figure 2:**
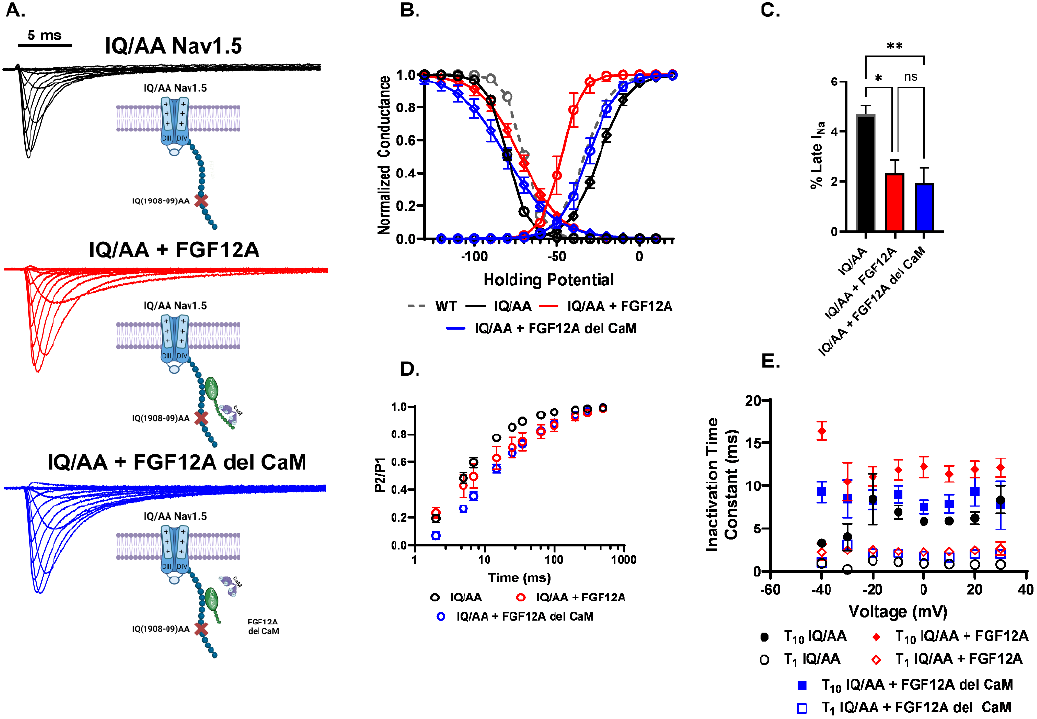
Removing CaM from the Nav1.5 CTD induces FGF12A-CaM dependent shifts in voltage-dependencies. **A)** Representative traces of the IQ(1908-09)AA Nav1.5 mutation which prevents binding of the CaM protein to the Nav1.5 CTD. Shown with schematic representations of the auxiliary subunits. **B)** Mean ±SEM GV and SSI plots with fitted Boltzmann functions. WT Nav1.5 (dashed grey), IQ/AA (black), IQ/AA+FGF12A (red), and IQ/AA+FGF12A del CaM (blue). The complete absence of CaM on the Nav1.5 CTD allows for a more dramatic CaM-dependent shift in activation and inactivation with the presence of WT FGF12A. **C)** Persistent late current (I_Na,L_) is significantly lowered regardless of the presence of CaM with FGF12A (Table 1). Percent of late I_Na_ at -30mV holding potential analysis is described in Methods. Error bars are SEM (* p = 0.0137, ** p = 0.0062). **D)** Mean ±SEM recovery from inactivation. **E)** Mean ±SEM T_1_ and T_10_ inactivation time constants of I_Na_ decay with WT FGF12A showing a significantly higher T_10_ component.

When co-transfected with the IQ/AA variant, a clear distinction in the SSI voltage-dependence was observed between WT FGF12A and FGF12A del CaM. Only WT FGF12A produced a significant depolarizing shift (+12.29mV ±1.787, p<0.0001) compared to the IQ/AA channel (**Figure 2B and Table 1**). A major hyperpolarizing shift in the GV curve was also observed in WT FGF12A (+15.07mV ±2.934, p<0.0001) with FGF12A del CaM shifting the GV V_1/2_ back to WT Nav1.5 values (−8.941mV ± 1.493, p<0.0001) (**Figure 2B and Table 1**). Similarly, only WT FGF12A significantly increased T_10_ (+6.074ms ±1.108, p=0.0009). However, both WT FGF12A and FGF12A del CaM slightly increased T_1_ (+1.55ms ±0.123, p<0.0001) and (+1.123ms ±0.281, p=0.0052) respectively compared to the IQ/AA variant (**Figure 2E and SI Table 1**). Accordingly, we discerned that in the absence of CaM already present on the Nav1.5 CTD, the addition of FGF12A drastically alters the inactivation gating of the Nav1.5 channel potentially through a CaM presence on the FGF12A N-terminal.

Conversely, the addition WT FGF12A blocks the increased levels of I_Na,L_ observed with the IQ/AA variant (−2.332% ±0.656, p=0.0137) as expected, however, the addition FGF12A del CaM also decreased the I_Na,L_ (−2.753% ±0.696, p=0.0062) (**Figure 2C**). The result of the reduction of pathogenic I_Na,L_ by either WT FGF12A or FGF12A del CaM led us to hypothesize that there are two different mechanisms by which FGF12A interacts with the Nav1.5 channel: CaM-regulated and CaM-independent.

### Impaired fast inactivation of Nav1.5 illustrates a CaM-independent mechanism of FGF12A

The F1486Q (IQM) mutation was used to inhibit fast inactivation of the Nav1.5 channel by impairing the inactivation gating particle located on the DIII-DIV linker (West et al., 1992). The marked difference in fast inactivation of the IQM variant is shown in **Figure 3A**. Additionally, IQM substantially increases I_Na,L_ compared to the WT Nav1.5 channels (+12.85% ±2.165, p<0.0001) (**Table 1**). The additions of either WT FGF12A (−6.832% ±2.16, p=0.0157) and FGF12A del CaM (−8.92% ±2.14, p=0.0396) were sufficient to reduce I_Na,L_ without changes to the peak current expression (**Figure 3D and Table 1**). Both WT FGF12A and FGF12A del CaM altered the time course of recovery from inactivation of the IQM variant channel in a similar manner (**Figure 3C**). GV and SSI curves in addition to fitted inactivation time constants for the IQM variant with(out) FGF12A are shown in Supplemental Information (**SI Figure 1 and 3**)

**Figure 3:**
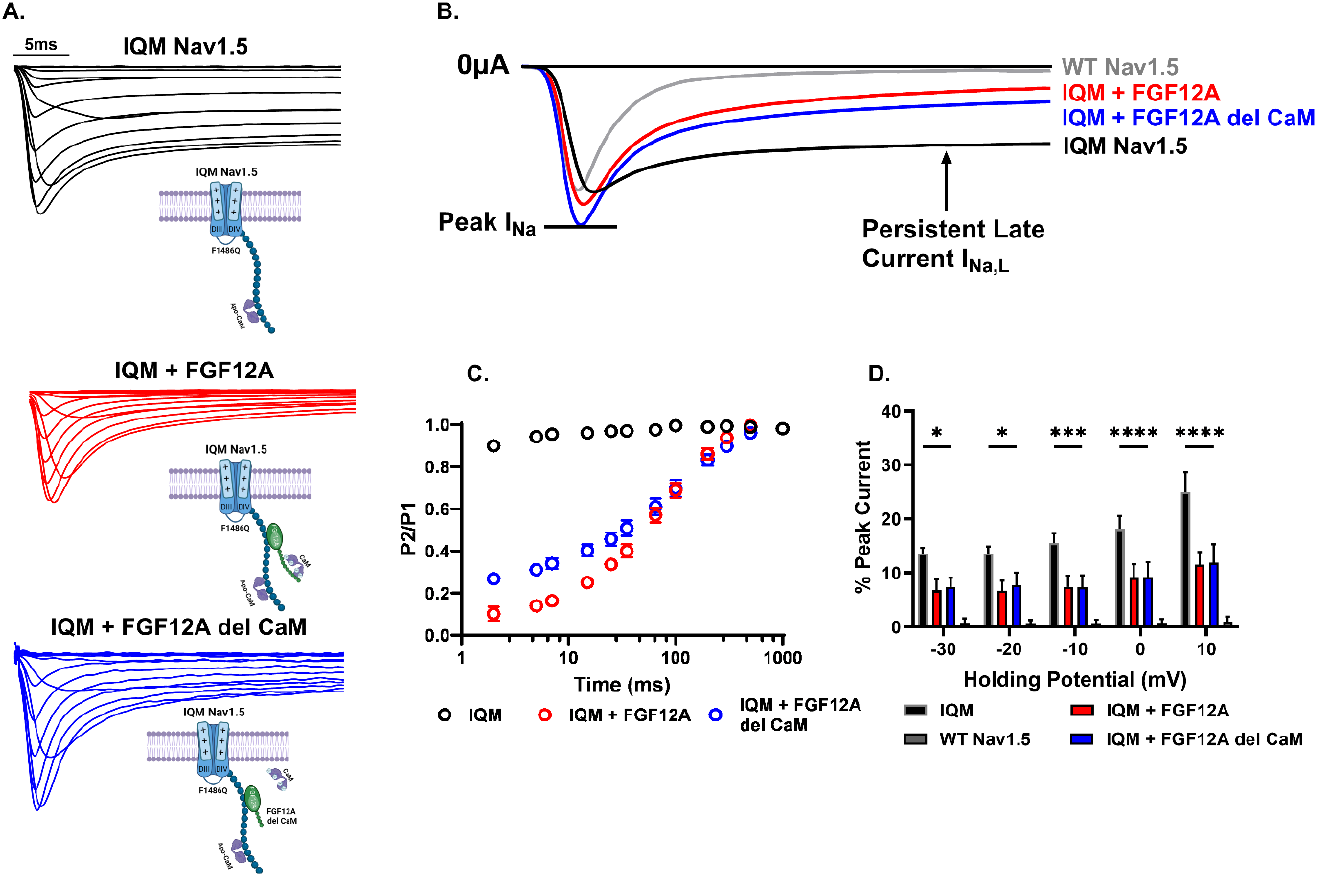
Inhibiting fast inactivation via the IQM mutation demonstrates the CaM-independent mechanism by which FGF12A inhibits I_Na,L_. **A)** Representative traces and schematic views of the F1486Q (IQM) variant, with(out) auxiliary subunits, that inhibits fast inactivation through altering the DIII-DIV linker responsible for fast inactivation after VSD DIV activation. **B)** Comparative traces of Nav1.5 at a step potential of -40mV. Comparison of persistent late current between Nav.15 variants includes determining the percentage of the Peak I_Na_ that the I_Na,L_ is. **C)** Recovery from inactivation shows that FGF12A with and without CaM significantly alters the relationship. **D)** Percentage of peak current comparing the differences in I_Na,L_ in WT and IQM channels at multiple holding potentials (mV). Error bars are ±SEM. (* p = 0.0396, 0.0234; *** p = 0.0004; **** p < 0.0001)

The reduction of I_Na,L_ in the IQM variant by both WT FGF12A and FGF12A del CaM is consistent with the results found in our experiments with the IQ/AA variant (**Figure 2C**). However, with the IQM variant, CaM is present in the Nav1.5 CTD, therefore we conclude that the process by which FGF12A reduces pathogenic I_Na,L_ is CaM-independent as the presence or absence of CaM does not affect I_Na,L_ inhibition.

### Nav1.5 LQT3 variants show a consistent split in CaM mechanisms

We continued observing FGF12A regulation of Nav1.5 voltage dependence by using LQT3 mutations located on the CTD (Y1795C and S1904L) (Bankston et al., 2007; Rivolta et al., 2001) (**Figure 4A**). Independently, Y1795C shifts the SSI V_1/2_ leftward compared to the WT Nav1.5 channel (−9.7mV ±1.628, p<0.0001) while S1904L shifts the GV V_1/2_ rightward (+11.61mV ±2.575, p=0.0041) (**Table 1**). A depolarizing (right) shift of the V_1/2_ of activation can lead to late channel activation while a hyperpolarizing (left) shift of inactivation increases the number of channels activated at higher membrane potentials. Both effects can potentially lead to a prolonged cardiac action potential and subsequent arrhythmias(Han et al., 2018).

**Figure 4:**
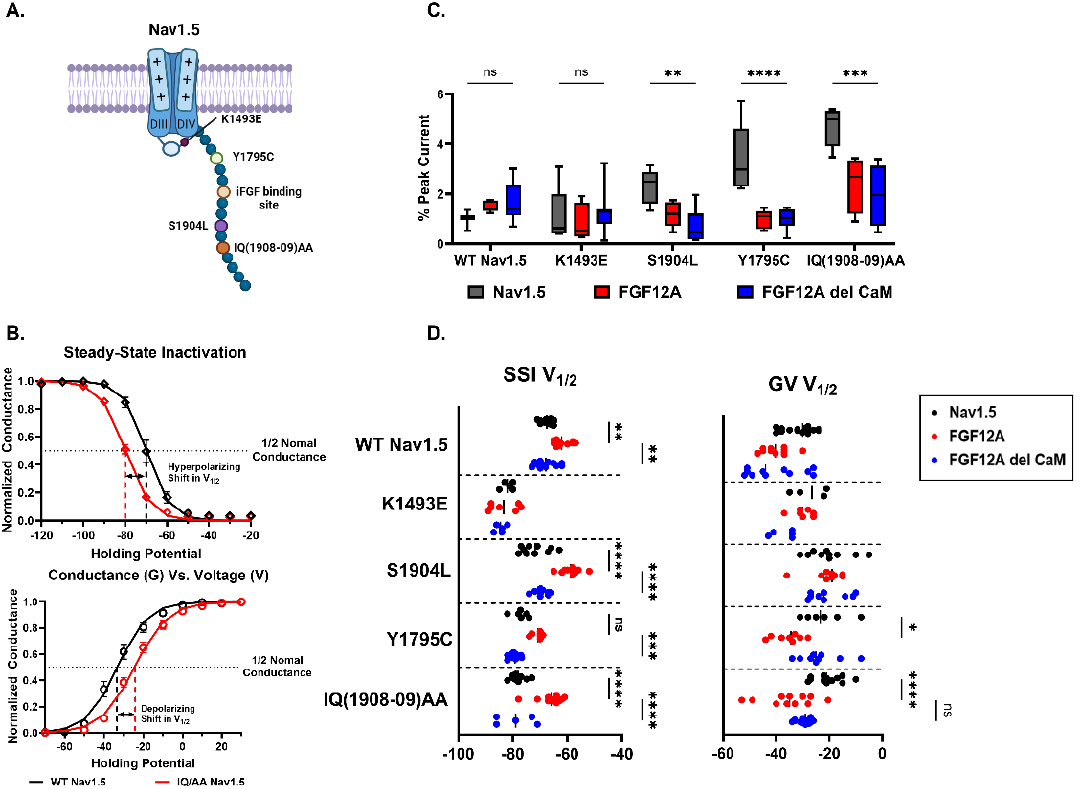
Altering Nav1.5’s gating kinetics show both CaM independent and dependent effects. **A)** Schematic representation of various Nav1.5 mutation locations tested. **B)** Example graphs of steady state inactivation (top) and conductance vs. voltage (bottom) that illustrate how the IQ/AA variant (red) shifts the V_1/2_ value in either a hyperpolarizing or depolarizing manner compared to WT Nav1.5 (black). **C)** FGF12A doesn’t require CaM to decrease the persistent late current from an altered Nav1.5 channel. Each Nav1.5 variant is separated into three groups: no FGF12A (grey), with WT FGF12A (red), with FGF12A del CaM (blue). Graph is box-and-whisker plot with bars from 5^th^ to 95^th^ percentile. (** p = 0.0058, *** p = 0.0002, **** p< 0.0001). **D)** FGF12A requires CaM to depolarize the SSI V_1/2_ except when the DIII-DIV linker is unable to interact with the CTD as in the case of K1493E. Additionally, a slight hyperpolarizing shift occurs with GV V_1/2_ only with WT FGF12A. Graph is a scatter dot plot with the median line shown. SSI (** p = 0.0032, 0.0019, *** p = 0.0003, **** p < 0.0001), GV (* p = 0.0289,0.0251, **** p < 0.0001) (**Table 1**).

We observed a trend in the SSI V_1/2_ with WT FGF12A depolarizing the V_1/2_ and FGF12A del CaM having no significant shifts in V_1/2_ ((**Figure 4D and Table 1**). We also observed that WT FGF12A induces a hyperpolarizing shift in the GV V_1/2_ of Y1795C, but not in S1904L (**Figure 4D and Table 1**). Thus, FGF12A alters the two variants in an opposite manner, depolarizing the SSI in S1904L and hyperpolarizing the GV in Y1795C. Additionally, the observation of WT FGF12A affecting SSI or GV while FGF12A del CaM does not is recapitulated with the IQ/AA variant (**Figure 2B**), which indicates that the mechanism is dependent on the CaM associated with FGF12A.

Conversely, this depolarizing SSI V_1/2_ trend was not shown in the K1493E variant. The K1493E variant individually shifts the SSI V_1/2_ to depolarized potentials (−14.3mV ±1.746, p<0.0001) compared to WT Nav1.5 (**Table 1**). The addition of either WT FGF12A or FGF12A del CaM does alter the already shifted SSI curve (**Figure 4D**). We hypothesize that this lack of effect is due to the requirement of Nav1.5 CTD interacting with the DIII-DIV linker which is impaired in the K1493E variant (Gade et al., 2019).

We continued to examine the FGF12A effect on the Nav1.5 I_Na,L_. S1904L and Y1795C both increased I_Na,L_ compared to WT Nav1.5 (+1.331% ±0.523, p= 0.0085) and (+2.431% ±0.562, p=0.0038) (**Table 1**). As predicted, the additions of either WT FGF12A or FGF12A del CaM were sufficient to reduce the pathogenic I_Na,L_ (**Figure 4C and Table 1**) consistent with our findings with the IQ/AA variant (**Figure 2C**) and the IQM variant (**Figure 3**). Representative traces and other measurements (GV, SSI, Recovery from Inactivation, and Inactivation time constants) are shown in the Supplemental Information Figures (**SI Figures 1-4)**. Overall, there was an established difference between the FGF12A ability to alter the voltage dependence of inactivation versus the persistent late current of the Nav1.5 channel with the former requiring CaM present on the FGF12A N-terminus.

### Intermediate inactivation indicates how FGF12A alters inactivation of the Nav1.5 channel

In addition to fast inactivation which occurs over a 1 to 10ms time domain, Na^+^ channels display inactivation over multiple slower time domains as well (Ruben et al., 1992). Intermediate inactivation is a distinct gating process that has a time period of 500ms to 1 second (Kambouris et al., 1998). As we have shown that FGF12A differentially alters the decay of fast inactivation and persistent late current, we proceeded to test the CaM effect on intermediate inactivation (**Figure 5**) utilizing protocols developed by Wang et al. 2000. We observed a hyperpolarizing shift in the intermediate inactivation between WT Nav1.5 and FGF12A del CaM (−6.856mV ±1.882, p=0.0015) (**Figure 5B**). WT FGF12A caused a non-significant depolarizing shift in the intermediate inactivation suggesting that only FGF12A without CaM present shifts the voltage-dependency of intermediate inactivation.

**Figure 5:**
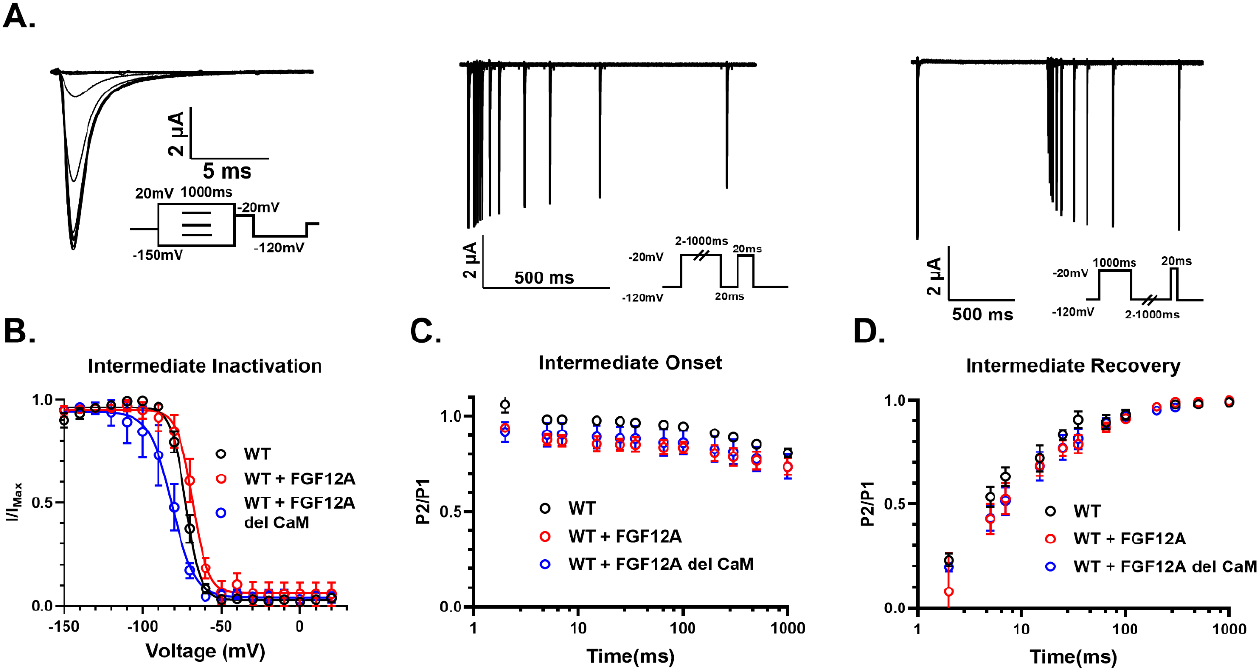
Intermediate inactivation of Nav1.5 shows the split mechanism between FGF12A and CaM. **A)** Representative trace and pulse protocols of (right to left): intermediate inactivation, intermediate onset, and intermediate recovery. **B)** Normalized ±SEM current vs voltage of intermediate with fitted Boltzmann function show WT (black, n=5), WT + FGF12A (red, n=4), and WT + FGF12A del CaM (blue, n=6). FGF12A requires CaM to function as a normal Nav1.5 channel. **C)** Normalized ±SEM values of the intermediate onset. FGF12A with and without CaM function similarly with a reduced onset compared to WT Nav1.5 without FGF12A. **D)** Normalized ±SEM values of intermediate recovery.

However, for the intermediate inactivation onset and recovery of the Nav1.5 channel, WT FGF12A and FGF12A del CaM had similar effects (**Figure 5C-D**). The reduced onset of intermediate inactivation due to either WT FGF12A or FGF12A del CaM implies a lower channel availability over a longer time period. The intermediate inactivation shifts could be associated with the reduction in persistent late current due to FGF12A as it has been shown with slow inactivation (Silva and Goldstein, 2013a, 2013c).

## Discussion

We have demonstrated the distinct differences in the CaM-independent and CaM-regulated mechanisms associated with FGF12A on the Nav1.5 channel. WT FGF12A when co-transfected with the Nav1.5 channel shifts the SSI in a depolarizing manner. This shift occurs only when CaM is present on FGF12A’s N-terminal. The effect is more pronounced when CaM is also removed from the Nav1.5 CTD. Conversely, when the fast inactivation of the channel is blocked via the IQM mutation, FGF12A, regardless of CaM’s presence, can rescue fast inactivation and reduce the persistent late current. The split mechanism trend was upheld when LQT3 variants were tested with the presence of WT FGF12A and FGF12A del CaM.

### The Nav1.5 CTD interacts with the DIII-DIV linker to stabilize fast inactivation

Current models of Nav1.5 fast inactivation include a CTD interaction with the III-IV linker (Biswas et al., 2024). Structures of Nav1.5 in the open, activated state suggest that the CTD initially interacts with the III-IV linker at the beginning of fast inactivation to stabilize the pore domain of the channel (Biswas et al., 2024; Shen et al., 2017). Once VSD-IV has activated and the IFM motif has tightly bound itself to the IFM receptor, the CTD moves away from the linker. Gade et al. 2019 altered the CTD’s interaction with the DIII-DIV linker with point mutations at K1493 and E1784 located on the DIII-DIV linker and CTD respectively (Gade et al., 2019). Changes in these residues provoked shifts in the fast inactivation of the channel and altered responses to pathogenic I_Na,L_ and indicate a reduction in interaction between the DIII-DIV linker and the CTD for normal channel conductance (Gade et al., 2019; Gardill et al., 2018). Increasingly, it has been shown that auxiliary subunits bound to the Nav1.5 CTD also interact with DIII-DIV linker/stabilize the overall interaction. CaM typically binds to the IQ motif on the Nav1.5 CTD (Kim et al., 2004), but has also been proposed to bind to a section of the DIII-DIV linker (Potet et al., 2009). Structural models have correspondingly shown both FGF12A and CaM located on the Nav1.5 CTD interacting with the DIII-DIV residues (Gade et al., 2019). CaM itself has been proposed to stabilize the interaction of the CTD with the IFM motif and to allow for faster state switching (Gabelli et al., 2014).

### FGF12A CaM-independent mechanism reduces pathogenic I_Na,L_

Previously published literature has touched on FGF12A’s effect of I_Na,L_ by considering its N-terminal section an inactivation particle (Abrams et al., 2020; Chakouri et al., 2022; Dover et al., 2010; Gade et al., 2019; Goldfarb, 2024; Wang et al., 2012). In Chakouri et al. 2022, the group proposed that the FGF12A N-terminal residues of 1-30 were sufficient to provide a protective effect against pathogenic I_Na,L_ including late current induced by the IQ/AA variant of Nav1.5 (Chakouri et al., 2022). Additionally, in Mahling et al. 2021, iFGF A-splice variants were proposed to have a long-term inactivation particle (LTP) on their N-terminus (Mahling et al., 2021). We hypothesize that FGF12A interacts with the DIII-DIV linker via its N-terminal (*residues 1-30*) to stabilize fast and slow inactivation. FGF12A can accomplish the same effects with the CaM-binding region deleted, negating the region’s necessity in reducing I_Na,L_. FGF12B, which is the splice variant of FGF12 without the N-terminal section of FGF12A, does not have the same reduction of I_Na,L_ as FGF12A (Chakouri et al., 2022). Goldfarb et al. 2024 describe a similar mechanism of canonical FGF12A interaction of specifically the initial N-terminal portion of the protein interacting with the IFM motif to stabilize the inactivation state of the Nav1.5 channel (Goldfarb, 2024).

### FGF12A CaM-regulated mechanisms alter inactivation kinetics of Nav1.5

The various iFGFs are prevalent throughout mammalian cells with FGF12 as the predominant isoform located within human cardiomyocytes andFGF13 as the most common in murine cardiomyocytes(Park et al., 2016; Wang et al., 2011). In Angsutararux et al. 2023, they determined that both the iFGF core region and the N-terminal of the protein is necessary to confer the shifts in inactivation and activation voltage-dependencies (Angsutararux et al., 2023). With regards to FGF12A, we showed that the presence of the CaM binding region altered FGF12A’s effects on both the steady-state inactivation and inactivation decay rate. FGF12B, which doesn’t have a CaM binding region on its N-terminus, can also shift both the SSI and GV in a depolarizing manner (Angsutararux et al., 2023). FGF12A independent of CaM does slightly alter the gating function of Nav1.5 as seen in **Figures 2 and 4**. However, FGF12A does require CaM for the full expected shifts in SSI and an increase the slow time component of inactivation. Potentially, FGF12A independently alters the VSD activations as seen in FGF12B (Angsutararux et al., 2023) whereas CaM interacts directly with the III-IV linker to alter Nav1.5 inactivation.

### Physiological Relevance

FGF12A has been shown to reduce pathogenic I_Na,L_ and shift the voltage-dependencies of steady-state inactivation in a depolarizing manner. Both effects, through either the CaM-independent or CaM-regulated mechanism, can reduce the duration of the cardiac action potential (Abriel, 2007; Han et al., 2018; Moreau et al., 2015, 2013). Typically, these effects are most prevalent when perturbations to the Nav1.5 channel have occurred either through genetic mutations such as LQT3 variants or changes in the presences of auxiliary subunits (Abriel, 2010; Han et al., 2018). FGF12A with respect to the persistent late current, could be viewed as a defense mechanism in restoring the normal action potential duration. With regards to the CaM present on the FGF12A N-terminal, the CaM binding site potentially could have evolved as an auxiliary protein redundancy to protect the fast inactivation of the Nav1.5 channel. The iFGF genes themselves are fairly conserved in all vertebrates with the CaM binding region being located on the N-terminal of all A-splice variants (Goldfarb, 2005; Satou et al., 2002). The IQ/AA Nav1.5 variant without CaM present shown increased I_Na,L_ and hyperpolarizing shifts in its SSI voltage dependencies. However, in IQ/AA transgenic mice, no effects are detected(Abrams et al., 2020). Abrams et al. 2020 proposed that the iFGFs endogenously present in the mice compensated for the lack of CaM on the Nav1.5 CTD (Abrams et al., 2020). However, it could be that the conserved CaM binding region on the A-splice variants of iFGFs are a secondary measure to ensure the presence of CaM on the Nav1.5 CTD.

### Overall FGF12A mechanisms with Nav1.5 gating states

As previously established by Mahling et al. 2021, all A-splice variants of iFGFs have a CaM binding region located on the N-terminal. Our results show that there are two distinct mechanisms by which FGF12A affects Nav1.5 conduction: a CaM-regulated and a CaM-independent mechanism. **Figure 6** represents these two mechanisms in a state dependent manner. At resting membrane potentials, the Nav1.5 channel is closed: all of its VSDs are in their downward positions and no Na^+^ ions are flowing (Chanda and Bezanilla, 2002; D et al., 2020). Once the membrane is depolarized, the VSD-I to III activate which translocates the coupled S5-S6 helices to open the pore, allowing for Na^+^ ions to flow inward (Chanda and Bezanilla, 2002; Jiang et al., 2021). Immediately after, VSD-IV activates revealing an allosteric pocket for the IFM motif allowing the III-IV linker to bind(Capes et al., 2013; Gardill et al., 2018; Varga et al., 2015). The binding forces the S5-S6 helices to constrict, causing fast inactivation of the channel, preventing the flow of Na^+^ ions (Biswas et al., 2024; Li et al., 2021). Between the activated and inactivated states of the channel, FGF12A and its associated CaM interact with the III-IV linker via the Nav1.5 CTD to alter the voltage dependencies of steady state inactivation and the fast inactivation decay rate (Angsutararux et al., 2023; Gade et al., 2019). We propose that instead of the FGF12A protein independently, the effects on Nav1.5 gating are due to the core of FGF12A bound to the Nav1.5 CTD (Angsutararux et al., 2023) and are regulated by the associated CaM protein. Once the Nav1.5 channel is inactivated, the FGF12A N-terminal will then interact with the DIII-DIV linker to stabilize the inactivated state of the channel and prevent pathogenic persistent late current consistent with previous publications (Chakouri et al., 2022; Dover et al., 2010; Goldfarb, 2024).

**Figure 6:**
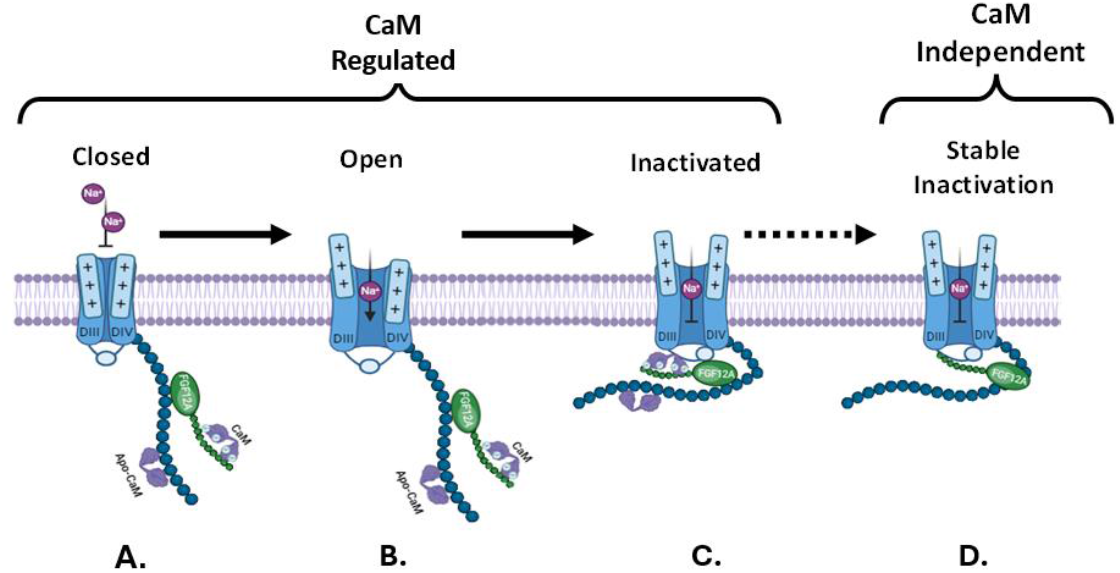
There are two mechanisms by which FGF12A interacts with the Nav1.5 channel via the CTD. **A)** <u>Closed state:</u> All VSDs are in their downward state; no Na^+^ ions are flowing inward. **B)** <u>Open state:</u> VSDs I-III are activated; Na^+^ ions are flowing inward. **C)** <u>Inactivated State:</u> The DIV VSD activates allowing the IFM motif to bind for fast inactivation to the allosteric pocket, thus shifted the coupled S5-S6 helices to close the pore; the Nav1.5 CTD swings to interact via CaM (on FGF12A) to modulate VSD activation. **D)** <u>Stably Inactivated State:</u> The N-terminal of FGF12A interact with the DIII-DIV linker to stabilize the inactivated state and prevent I_Na,L_.

---

**Supplemental Table.**

| Variant | Decay inactivation (n) | T<sub>1</sub> | ±SEM | T<sub>10</sub> | ±SEM |
| --- | --- | --- | --- | --- | --- |
| Nav1.5 | 4 | 2.776 | 0.3791 | 3.926 | 1.335 |
| Nav1.5 + FGF12A | 5 | 1.504 | 0.1669 | 9.917 | 0.7506 |
| Nav1.5 + FGF12A del CaM | 4 | 1.204 | 0.1454 | 7.203 | 1.072 |
| Nav1.5 + FGF12A Scram CaM | 5 | 1.997 | 0.7379 | 5.535 | 1.505 |
| IQ/AA | 3 | 0.2276 | 0.0031 | 4.008 | 1.565 |
| IQ/AA + FGF12A | 4 | 2.456 | 0.3141 | 10.5 | 2.183 |
| IQ/AA + FGF12A del CaM | 7 | 3.104 | 0.3358 | 8.493 | 2.243 |
| IQM | 7 | 3.644 | 0.2744 | 17.2 | 3.67 |
| IQM + FGF12A | 7 | 3.01 | 0.2447 | 20.99 | 1.245 |
| IQM + FGF12A del CaM | 7 | 1.885 | 0.1275 | 15.93 | 1.364 |
| K1493E | 5 | 0.9726 | 0.09687 | 5.605 | 0.435 |
| K1493E + FGF12A | 6 | 0.939 | 0.1569 | 7.565 | 1.786 |
| K1493E + FGF12A del CaM | 7 | 1.357 | 0.2375 | 3.573 | 0.1386 |
| S1904L | 8 | 4.543 | 1.449 | 13.89 | 3.776 |
| S1904L + FGF12A | 7 | 1.559 | 0.5945 | 16.19 | 5.726 |
| S1904L + FGF12A del CaM | 5 | 4.083 | 1.244 | 13 | 4.734 |
| Y1795C | 5 | 1.685 | 0.4691 | 4.964 | 1.44 |
| Y1795C + FGF12A | 8 | 4.862 | 0.1627 | 16.69 | 4.242 |
| Y1795C + FGF12A del CaM | 8 | 1.9999 | 0.353 | 6.039 | 1.725 |

## Supplemental Figures

**SI Figure 1:**
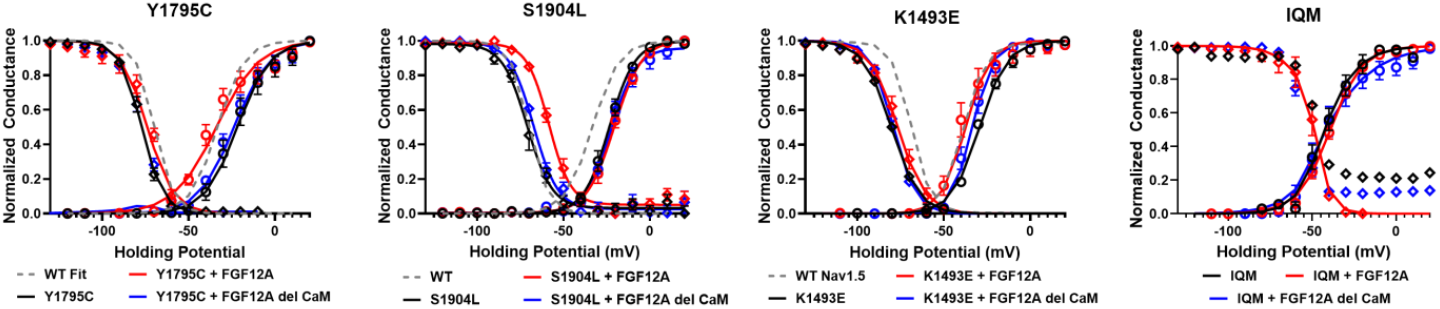
Mean normalized ±SEM conductance-voltage (GV) and steady-state inactivation (SSI) curves for WT Nav1.5 and the addition of: FGF12A, FGF12A del CaM, and FGF12A Scram CaM. Curves are fitted single Boltzmann functions.V_1/2_ values are recorded in **Figure 4** and **Table 1**. Shifts in the SSI and GV are seen withthe addition of only WT FGF12A in the S1904L and Y1795C variants respectively, requiring CaM for the full effects. The K1493E variant inhibits intercation of the Nav1.5 CTD with the III-IV linker, therefore no shifts are observed with the addition of WT FGF12A or FGF12A del CaM. Theaddition of FGF12A to the IQM mutation rescued fast inactivation without shiftingthe voltage-dependency of inactivation.

**SI Figure 2:**
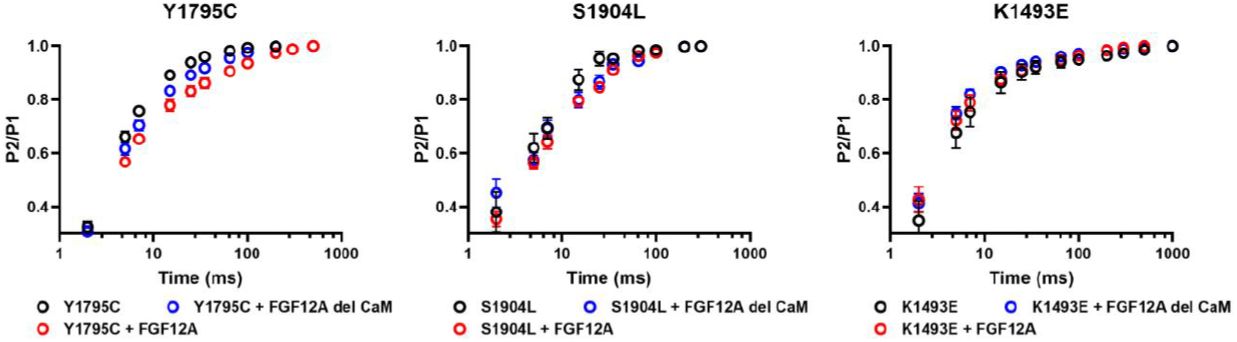
Mean ±SEM recovery from inactivation of the Y1795C, S1904L, and K1493E variants. No significant shifts were detected.

**SI Figure 3:**
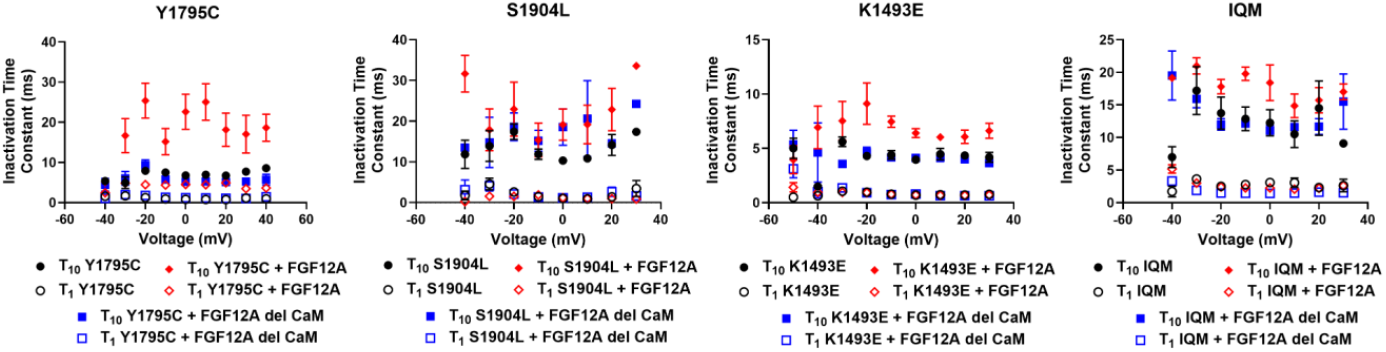
Mean ±SEM _1_ and _10_ inactivation time constantsof peak I_Na_ decay with the addition of WT FGF12A with the Y1795C, S1904L, K1493E, and IQM variants showinga significantly higher _10_ component.Values at holding voltage of-30mV are show in **SI Table 1**.

**SI Figure 4:**
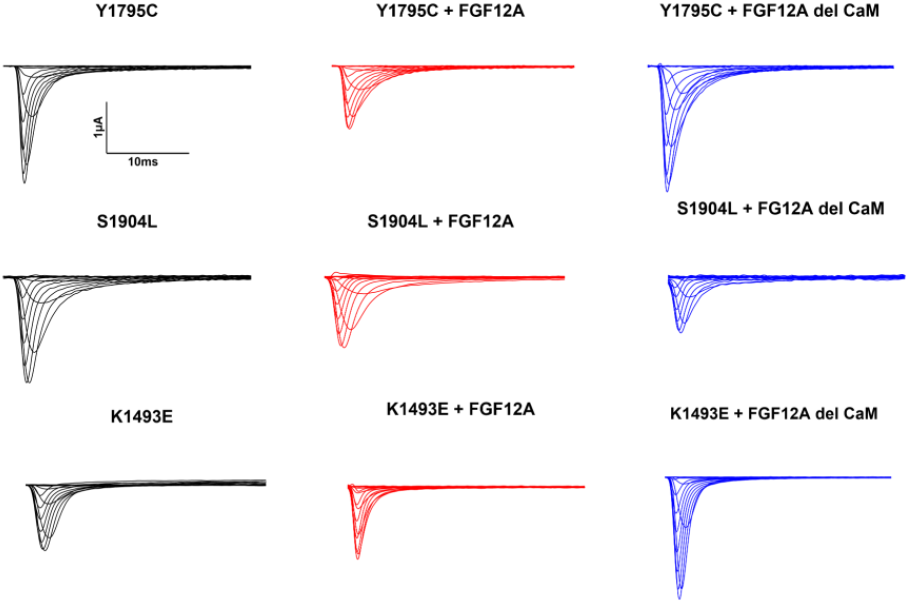
Representative traces of variants shown in **Figure 4**. Current traces were filtered via bandpass filter in Clampfit and are not normalized.

## Acknowledgements

This work was completed in its entirety within the Silva lab at Washington University in St. Louis. It was funded by NIH-NHLBI RO1 HL136553, NIH-NHLBI RO1HL150637 and the individual predoctoral fellowship 1F31HL164080-01A1.

